# Scaling up biodiversity ecosystem functioning relationships: the role of environmental heterogeneity in space and time

**DOI:** 10.1101/2020.11.04.367250

**Authors:** Patrick L. Thompson, Sonia Kéfi, Yuval R. Zelnik, Laura E. Dee, Shaopeng Wang, Claire de Mazancourt, Michel Loreau, Andrew Gonzalez

## Abstract

The biodiversity and ecosystem functioning (BEF) relationship is expected to be scale-dependent. The autocorrelation of environmental heterogeneity is hypothesized to explain this scale dependence because it influences how quickly biodiversity accumulates over space or time. However, this link has yet to be demonstrated in a formal model. Here we use a Lotka-Volterra competition model to simulate community dynamics when environmental conditions vary across either space or time. Species differ in their optimal environmental conditions, which results in turnover in community composition. We vary biodiversity by modelling communities with different sized regional species pools and ask how the amount of biomass per unit area depends on the number of species present, and the spatial or temporal scale at which it is measured. We find that more biodiversity is required to maintain functioning at larger temporal and spatial scales. The number of species required increases quickly when environmental autocorrelation is low, and slowly when autocorrelation is high. Both spatial and temporal environmental heterogeneity led to scale dependence in BEF, but autocorrelation had larger impacts when environmental change was temporal. These findings show how the biodiversity required to maintain functioning is expected to increase over space and time.

## Introduction

Biodiversity change arising from the loss and gain of species is now understood to affect many ecosystem processes, such as primary and secondary production, that together define how ecosystems function [1]. To date, our understanding of how biodiversity change affects ecosystem functioning (i.e. the so-called Biodiversity Ecosystem Functioning relationship, or BEF) is founded on tests of theory with controlled experiments and field observations conducted at relatively small scales of space and time [2]. However, recent research has found that the effects of biodiversity change on ecosystem functioning are scale dependent [3–5]. As a consequence, the slope of the BEF relationship (i.e., the rate at which function changes with biodiversity) will depend on the spatial and temporal scales over which it is measured. This scale dependence is important to consider for understanding the effects of biodiversity change on the functioning of ecosystems and on the services they provide to society [6].

Early research, conducted primarily on small spatial scales and over short durations, identified two ways in which biodiversity affects ecosystem functioning—complementarity effects and selection effects [7]. Complementarity effects emerge from niche differentiation and facilitation among species and lead to improved performance of species assemblages, compared to what would be expected based on the performance of monocultures of the component species. Selection effects arise when the species that perform well in monoculture also come to dominate (i.e. have high abundance) mixed species assemblages. These two effects are known to drive ecosystem functioning to have a positive but decelerating relationship with increasing diversity within assemblages [8]. An important open question is how these mechanisms support the emergence of BEF relationships across ecological communities spanning larger and longer spatial and temporal scales [2].

Long-term experiments are revealing that the slope of the BEF relationship, can vary with environmental context across sites and through time [5,9–12]. Indeed, the longer the length of the interval of time or space, the more likely we are to find new species contributing to ecosystem processes in different environments. This effect is largely driven by environmental heterogeneity (the degree of dissimilarity in environmental conditions, e.g. temperature, pH, nutrients) among sites, or between time points, which sets the scale of species turnover (i.e. beta diversity) [13–15]. Compositional turnover in response to environmental heterogeneity can be considered a form of complementarity, when different species contribute to ecosystem functioning in different conditions [16]. Turnover in species composition across space, or over time, due to environmental heterogeneity is predicted to lead to BEF relationships at large spatial scales or long temporal scales that diverge from the BEF relationships that are observed at small and short scales [3]. Factors mediating the rate of community turnover in the distribution of species are therefore predicted to determine how the BEF relationship varies with scale.

Despite the importance of environmental heterogeneity in determining community turnover, to date, relatively little theory has assessed whether environmental heterogeneity can explain scale dependence in the BEF relationship [2,17]. We address this important gap herein. Previous theory identified how community turnover could make the BEF relationship scale dependent [3], but did not provide a mechanistic link between community turnover and environmental heterogeneity. Spatial turnover arises because species’ distributions are restricted and patchy. Factors explaining the patchiness of species distributions and biomass include dispersal limitation and environmental heterogeneity [18,19]. In turn, patchiness in species’ occupancy and productivity will govern where and when different species contribute to ecosystem functioning. Likewise, temporal turnover arises through colonization and extirpation dynamics that are driven by temporal environmental change, interactions between species, and dispersal [20]

Environmental autocorrelation describes the rate of decay in environmental similarity in time or space [16]. Low (high) autocorrelation defines high (low) rates of change in environmental conditions over short durations or distances resulting in rapid (slow) decay in environmental similarity. Environmental autocorrelation affects many properties of communities and ecosystems: Vasseur and Yodzis [16] found environmental time series to be autocorrelated, with aquatic environments more so than terrestrial environments. Theory and experiments have demonstrated that population extinction probabilities [21,22] as well as many aspects of population [23–25] and community structure [26] and dynamics [15,27,28] are strongly modulated by environmental autocorrelation. Similarly, the effects of biodiversity on the mean and temporal variability of ecosystem functioning (e.g. biomass production) depend markedly on the degree of autocorrelation of environmental conditions because it sets the rate of species turnover [15,29,30]. However, this research did not address how environmental autocorrelation affects BEF relationships at different spatial and temporal scales.

Here we ask how environmental heterogeneity and its autocorrelation in space and time cause the BEF relationship to change with the spatial or temporal scale at which it is observed. We tested three hypotheses: H1 - the number of species required to maintain ecosystem functioning will increase with spatial scale because species vary in the conditions in which they are productive. H2 - the degree of scale dependence in BEF will depend on the degree to which environmental autocorrelation (i.e., high vs. low autocorrelation) mediates turnover in species composition. H3 - the degree of scale dependence in BEF will depend on whether environmental autocorrelation is temporal or spatial. This is because, in the absence of strong dispersal, the composition of a community is more likely to depend on the previous conditions experienced rather than on the conditions in adjacent habitats [20]. We addressed these hypotheses by simulating landscapes of many communities with Lotka-Volterra dynamics driven by environmental heterogeneity in space and time, and where autocorrelation is controlled.

## Methods

We used numerical simulations of Lotka-Volterra competitive communities to explore how the relationship between biodiversity (as species richness) and ecosystem functioning (as total community biomass), hereafter the BEF relationship, depends on the spatial or temporal scale at which it is measured (Fig. S1).

We used two different scenarios to explore BEF scaling in time and space:

1. a *spatially variable scenario* of 80 patches arranged along a spatial environmental gradient, with no temporal change in environmental conditions.
2. a *temporally variable scenario* of one patch that experiences temporal fluctuations in environmental conditions over 80 time steps.

We chose 80 as our maximum spatial or temporal scale of interest because it is large enough to incorporate substantial environmental heterogeneity while remaining computationally tractable. Changing this maximal scale has little impact on our results (Fig. S3, S4) as long as this scale is sufficient to allow for environmental heterogeneity to result in variation in community composition.

### Environmental heterogeneity

In each scenario, we contrasted three levels of autocorrelation in environmental heterogeneity, low medium, and high. We define these levels of autocorrelation based on the inverse power law 1/*f*^γ^ [31] where γ corresponds to the level of autocorrelation (low = 0, medium = 1, and high = 2; Fig. S1D). Sequences of environmental heterogeneity corresponding to these three levels of autocorrelation were generated using the *phase.partnered* function in the *synchrony R* package [32]. These sequences defined how environmental conditions varied across space in the spatial scenarios and across time in the temporal scenarios. All three levels of autocorrelation had a mean value of 0.5 and a standard deviation of 0.25, so that local conditions, as defined by *env_x_*(*t*), are approximately in the range of 0 to 1.

### Species responses to environmental heterogeneity

We considered a global pool of 100 species which differed in their environmental optima *zi*. These optima were evenly distributed between −0.2 and 1.2, which covers the range of optima where species could have positive growth for any environmental condition included in our environmental sequences. The match between this environmental optimum and local conditions, *env_x_*(*t*), of the patch *x* in which species *i* is present, determines its density-independent rate of growth, *ri_x_*(*t*):

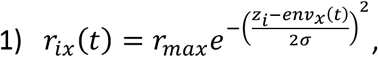

1) Where *σ* is equal to 0.25, and *r_max_* = 5 is the maximum density-independent growth rate, which occurs when the local environmental conditions match the species environmental optimum, i.e. *z_i_* = *env_x_*(*t*).

### Community dynamics

The temporal dynamics of the abundance of each species depend on both the density-independent rate of growth (eq. 1) and density-dependent competition between species [following 20]:

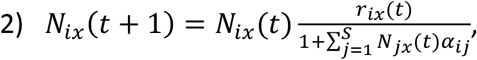

Where *N_ix_*(*t*) is the biomass of species *i* in patch *x* at time *t*. The per capita competition coefficients, *α_ij_*, determine the strength of intraspecific *α_ii_* and interspecific *α_ij_* densitydependent competition. We set all values of intraspecific competition *α_ii_* to 1 and draw values of *α_ij_* from a uniform distribution between 0 and 0.25. An important assumption is that intraspecific competition is stronger than interspecific competition. Without this assumption, a positive local BEF relationship would not emerge [20,33]. Because we assume that all species have equal r_max and equal intraspecific competition *α_ii_*, all species have equal carrying capacity when experiencing optimal environmental conditions. However, our sensitivity analyses show that our results are robust to variation in intraspecific competition (Appendix 2). We assume that there is no dispersal between patches, but that there is dispersal from an external species pool (see Supplementary materials, Appendix 1 - *Rationale for decisions in simulation*).

### Environmental sequences

We used the exact same 80 step environmental sequences for both our spatial and temporal scenarios. We ran the spatial simulations for 150 time steps, in order to allow the communities to reach equilibrium and then based our analysis on the community composition in the final time step. For the temporal simulations, we preceded the sampled environmental sequences with an 80 time step burn-in sequence with the same level of autocorrelation. This burn in sequence was not used in our analysis but allows the simulations to reach a dynamic equilibrium (i.e., no temporal trend in the average species richness and community biomass; Fig S2). For details on this burn-in sequence see the *Generating the temporal burn-in sequence* section in Appendix 1.

### Simulations

We considered three different levels of environmental autocorrelation for each of the spatially and temporally varying scenarios. This resulted in a total of six environmental sequence types. For each environmental sequence type, 100 different randomizations of the sequence and species interactions *α_ij_* were considered. For each individual environmental sequence, we ran the simulation with potential levels of species richness from 1 to 100 species. We did this by selecting a random subset, *S\**, of the full 100 potential species corresponding to the level of potential biodiversity. Each patch was initialized with all *S\** species, each with a biomass of 1. By running each environmental sequence with different numbers of species seeded into the simulation, *S\**, we were able to contrast different levels of species richness and thus determine the relationship between biodiversity and community biomass [*sensu* 39,40]. After initialization, we ran the simulation for *T* time steps, including all burn-in and sampled time steps (T = 150, 149 burn-in and 1 sampled time step in spatial scenario; T = 160, 80 burn in and 80 sampled for temporal scenario) for spatial (temporal), performing the following actions in each time step:

1. For each patch *x* and species *i*, update the biomass according to equation 2 (given the environment at time *t*).
2. If in any patch a species from the seeded species pool, *S\**, has a biomass value lower than 0.05, we consider it to be lost from that patch and set its biomass to zero.
3. Record the identity of all species that are present in each patch as well as their summed biomass.
4. Reintroduce all lost species in all patches, by setting their biomass values to 0.035 (below our extinction threshold).

Update the time (*t*= *t* +1), and go back to step 1; continue until the final time step is reached.

For a description of the rationale of the specific decisions in the model and the simulations, see supplementary materials (Appendix 1).

### Estimating the BEF relationship at multiple scales

We refer to scale as the number of time steps (in the temporally variable scenario) or the number of patches (in the spatially variable scenario) used to evaluate the BEF relationship. In each simulation, we calculated the cumulative biomass and species richness at all scales, from one patch to all 80 patches when examining scaling in space, or from one time step to 80 time steps when exploring scaling in time. We did this by aggregating patches in space or across time steps sequentially to increase the grain, or ‘scale’. We, for example, aggregate five sequential time steps or patches to evaluate the BEF at a scale of 5 and calculate species richness and the cumulative biomass (see Fig. S1). The sequential aggregation ensured that we maintained the spatial structure of the environment as we combined patches or time points. In all cases, we start at the final time step or patch and aggregate previous or adjacent time steps and patches. We performed this aggregation at all spatial and temporal scales from one to 30 patches (times), every two scale grains from 32 to 40, and then every 5 scale grains from 45 to 80. Then, at each spatial or temporal scale, we contrasted values of biomass and species richness from simulations with different species pool sizes but using the same sequence of environmental heterogeneity. We then used a Michaelis-Menten function [34] to estimate the relationship between species richness and average cumulative biomass (Fig. S1B): 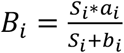, where *B_i_* is the total biomass at scale *i*, *S_i_* is the species richness, *a_i_* is saturation level for biomass at scale *i*, and *b_i_* is the number of species required to sustain half that asymptotic level of biomass (see dotted lines on Fig. S1B). Higher levels of *b* indicate that a greater number of species are required to maintain a given amount of biomass (e.g. yellow curve on Fig. S1B). To make it possible to determine whether biomass is accumulating linearly across space and time, we used the average biomass across all patches or time points considered. This averaging changed the asymptote of the BEF relationship that we observed but did not change the estimated number of species required to maintain biodiversity (i.e. the half saturation richness). Note that this is different from Thompson et al. [3] who estimated the slope of the BEF relationship (the rate at which log biomass increases per log species) instead of the half saturation richness. All simulations and analyses were performed in *R* v.3.6.1 [35] (available at - https://doi.org/10.5281/zenodo.4174454).

The critical tests of our hypotheses are thus: H1 – that the half saturation constant *b* of the BEF relationship increases with spatial or temporal scale. H2 – that the degree to which *b* changes with scale depends on the autocorrelation of the environment. H3 – that *b* increases with scale at a different rate when environmental change is temporal vs. spatial.

## Results

### Compositional turnover and species richness

The degree of environmental autocorrelation in either space or time determines the rate of compositional turnover. Compositional turnover, between neighbouring patches or time points, is high when environmental heterogeneity is uncorrelated (γ = 0) and decreases as the environmental autocorrelation increases (Fig. 1A-C for space; Fig. 1D-F for time). This compositional turnover leads to an increasing but saturating relationship between species richness and scale (Fig. 2A,B). Species richness increases and saturates fastest with scale when environmental heterogeneity is uncorrelated (γ = 0) because the full range of environmental conditions is encountered over short scales of space or time. These increases in species richness with scale are slower in autocorrelated environments (γ > 0) because short scales tend to only include a subset of environmental conditions. Species richness saturates at a slightly lower level when autocorrelation is high (γ = 2), because these environmental sequences tend to include fewer extreme values compared to when autocorrelation is lower. Species richness increases faster with spatial scale (Fig. 2A) compared to temporal scale (Fig. 2B) because communities are at static equilibrium with respect to the environment in the spatial scenario but not in the temporal environment. This non-equilibrium state in temporally changing environments results in a lag in how quickly species richness accumulates.

**Figure 1:**
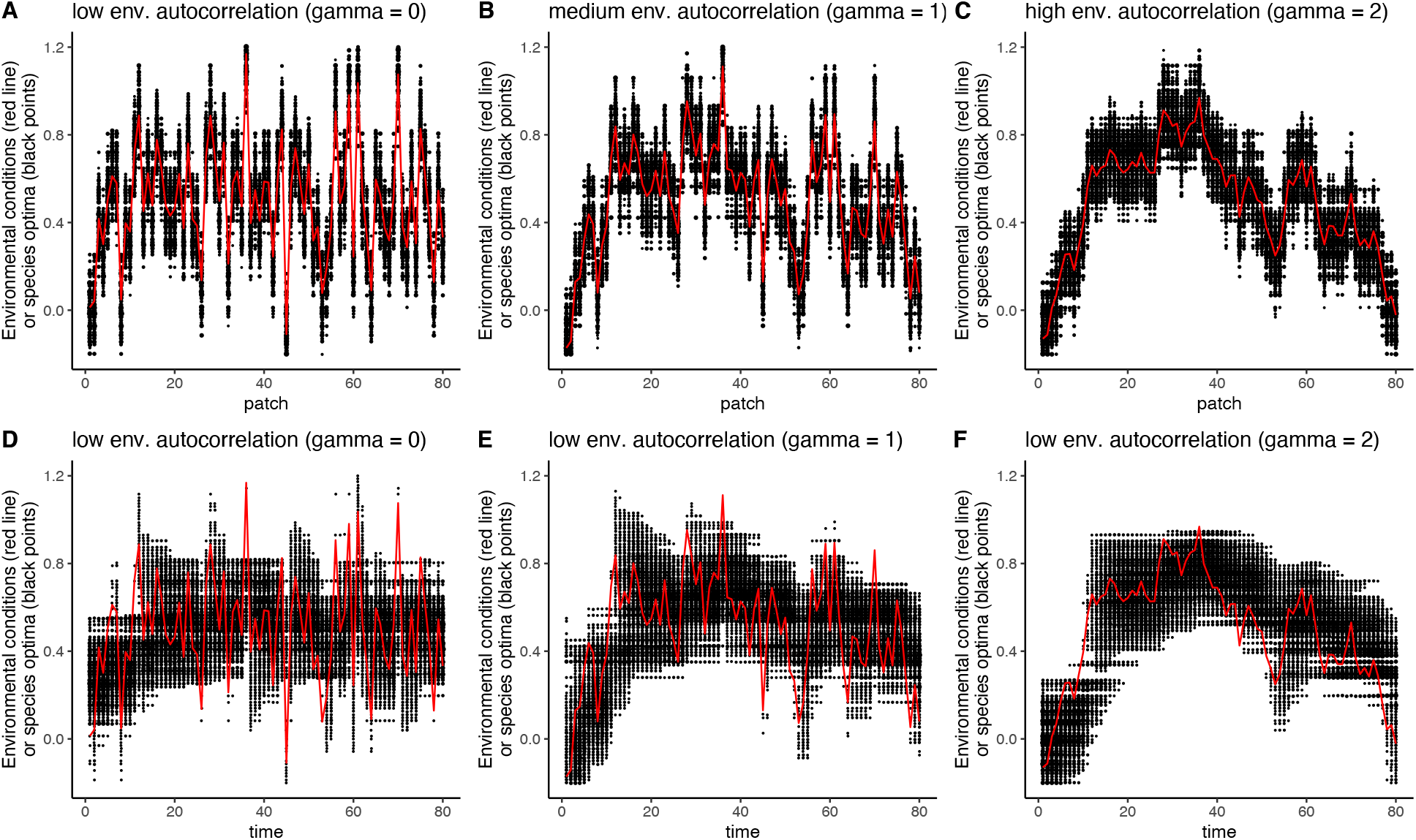
Spatial (A-C) and temporal (D-F) turnover in community composition as a function of environmental autocorrelation (A, D, low; B, E, medium; C, F, high). The red line shows the environmental value in a given patch (A-C) or time point (D-F), while the black points show the species that are present (positioned on the y-axis according to their environmental optima). Results from a single representative run of the model, with 100 potential species.

**Figure 2.**
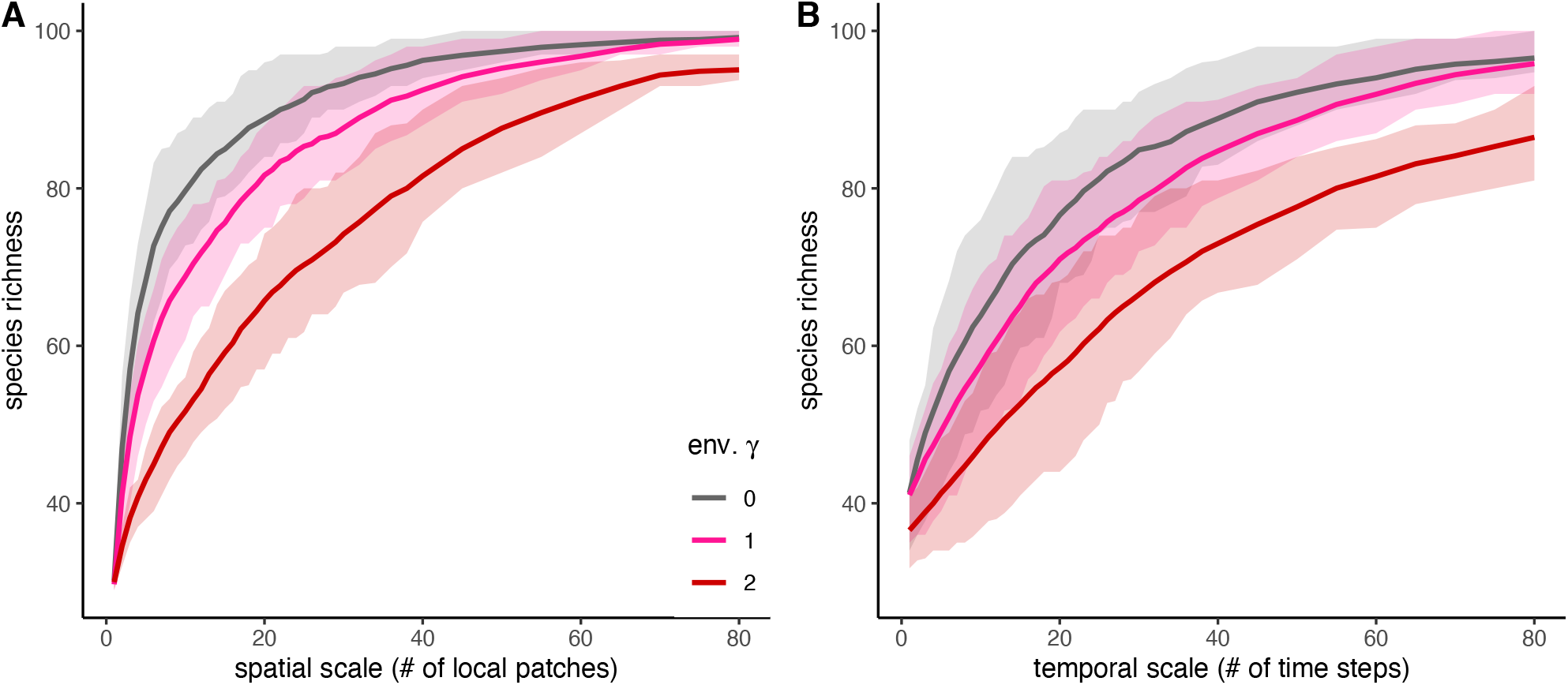
Species richness versus spatial scale (A), and temporal scale (B). Colour indicates the γ value of the degree of autocorrelation of environmental heterogeneity. Lines represent the mean and bands represent the interquartile range across 100 replicate simulations with the full species pool.

### Cumulative biomass by scale

Biomass accumulates with scale additively and so the average biomass per patch or time point does not depend on the number of patches or time points considered. Because communities are independent in space, the average biomass does not depend on how environmental conditions are autocorrelated across space (Fig. 3A-C). However, the degree of temporal autocorrelation does influence the average biomass that is produced (Fig. 3D-F). Biomass is higher when environmental conditions are temporally autocorrelated because species sequential environmental conditions allow species to establish and increase in abundance.

**Figure 3.**
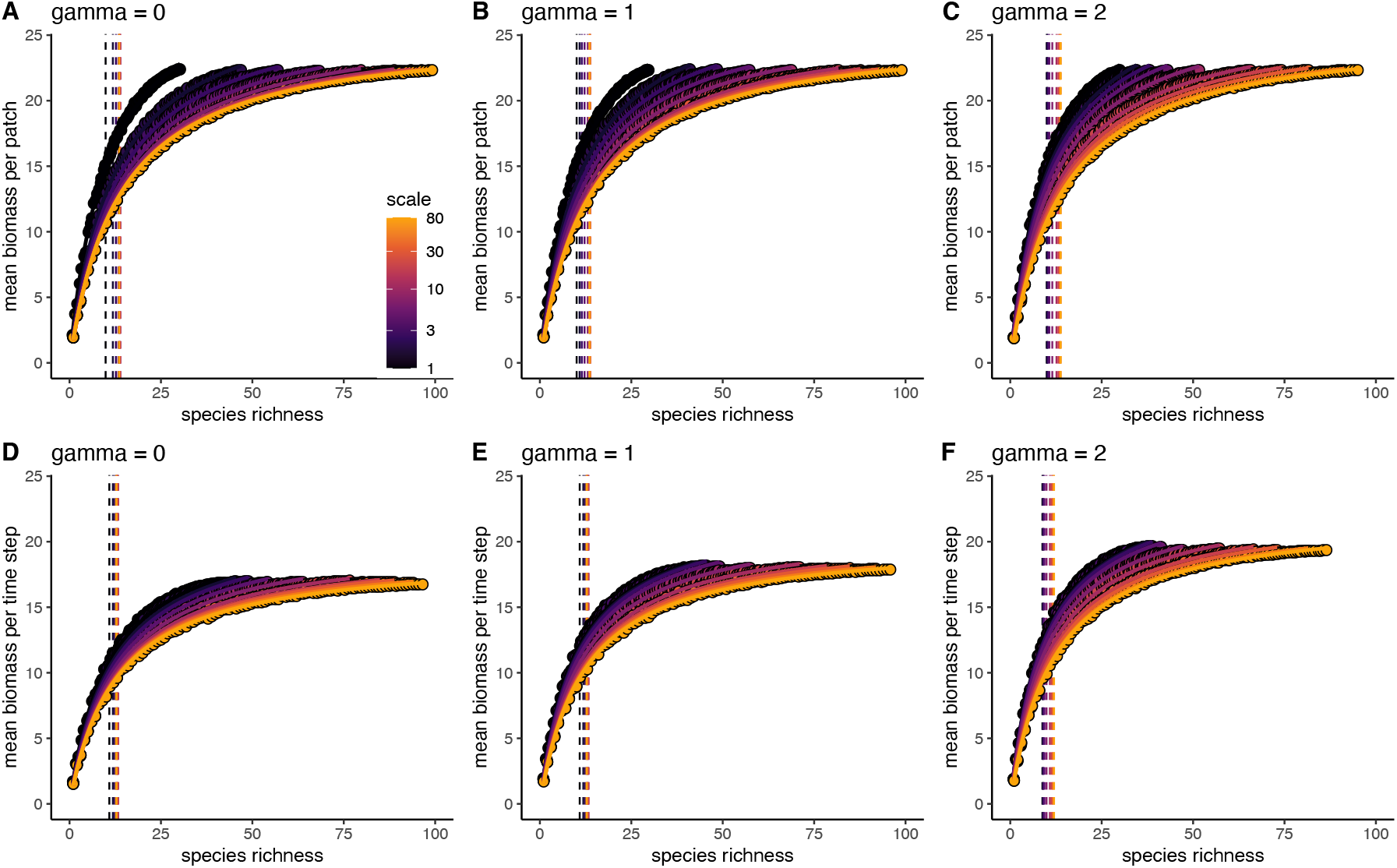
The relationship between species richness and mean biomass across space (A-C) or time (D-F) at multiple scales (colours) with different levels of environmental autocorrelation. To make the BEF curves visually comparable we use biomass per unit area (or unit of time). Environmental autocorrelation is different between panels (A, D, low (γ = 0); B, E, medium (γ = 1); C, F, high (γ = 2)). Colour indicates the scale at which the BEF relationship is observed at (darker = smaller or shorter, brighter = larger or longer). Variation in observed species richness and average biomass within a spatial scale results from different numbers of species used in the simulations (i.e. each data point is the value from a single simulation run with a different number of initial species *S*\*). Vertical lines show the mean value of the half saturation constant, estimated separately at each scale of observation. Data based on 100 replicate simulation runs.

### The biodiversity-ecosystem functioning relationship and scale

The half BEF saturation richness *b_i_* increases with both spatial and temporal scales (Fig. 3, 4). This means that more species are required to sustain the same amount of biomass per unit area or time over larger scales compared to smaller scales.

**Figure 4.**
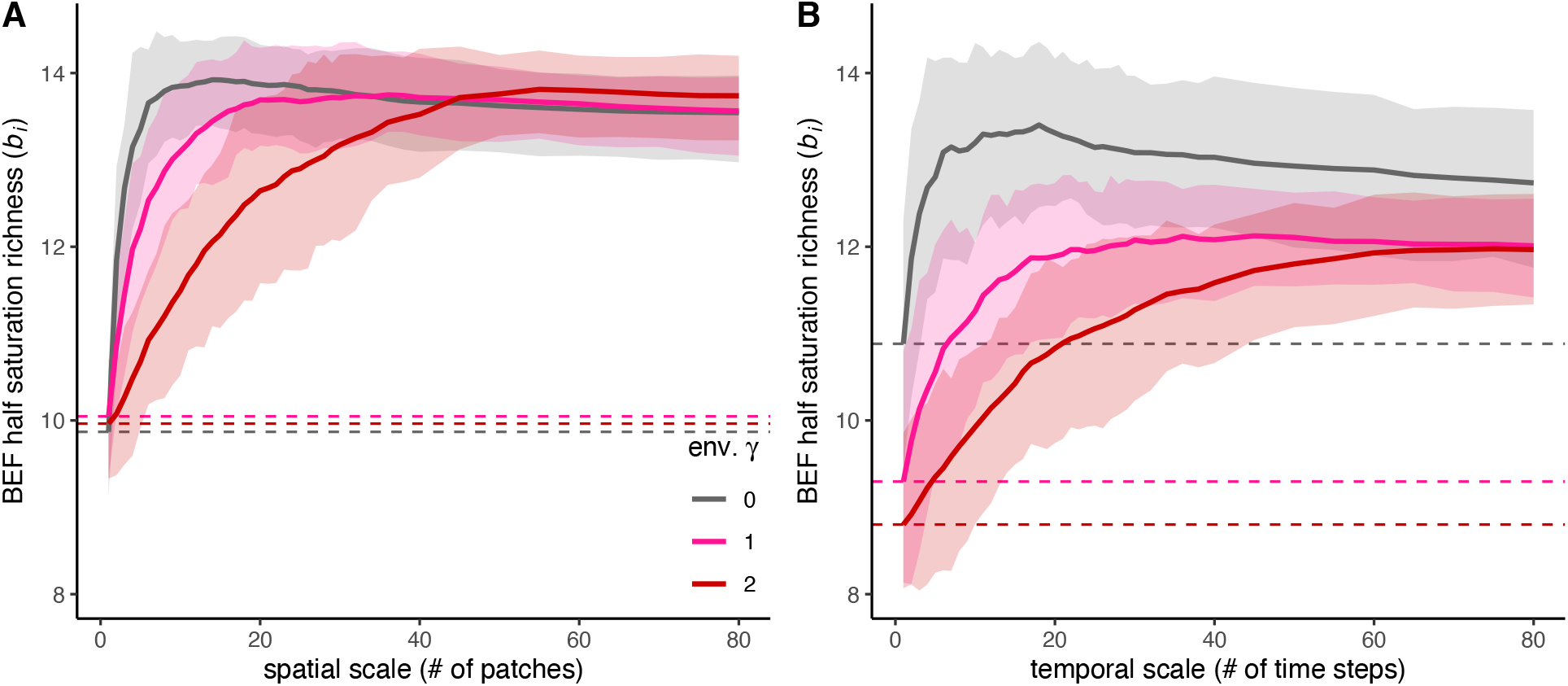
The relationship between the half saturation species richness of the BEF relationship and the spatial (A) or temporal (B) scale of observation. Colour indicates the γ value of the degree of autocorrelation of environmental heterogeneity. Dashed lines show the half saturation species richness at a scale of 1. Lines represent the mean and bands represent the interquartile range across 100 replicate simulations.

When environmental conditions vary across space, the degree of autocorrelation has no effect on community composition or biomass at local scales, and so *b_1_* (at the single patch scale) has nearly the same value (approx. 8.5; see Fig. 4A) in all cases. But the rate at which *b_i_* increases with scale depends on the degree of environmental autocorrelation, with the fastest increases when environmental autocorrelation is low (i.e. fastest when γ = 0, slowest when γ = 2). This is because more species are required to maintain productivity under the high heterogeneity encountered over small spatial scales when the environment varies randomly. But over larger spatial scales, a greater range of environmental conditions is encountered, regardless of the degree of environmental autocorrelation, and so *b_i_* saturates at roughly the same level (approx. 13.5; see Fig. 4A) in all cases. This result is highly dependent on our assumption that a similar range of environmental conditions experienced over large or long scales is the same, regardless of the degree of autocorrelation.

When environmental heterogeneity is across time, it impacts communities at short time scales. The local scale *b_i_* is greatest in randomly varying environments (*b_1_* = 9.3 when γ = 0) and lowest in environments with high autocorrelation (*b_1_* = 8.1 when γ = 2; Fig. 4). This is because overall biomass is reduced by random temporal environmental heterogeneity (Fig. 3) since species never experience sustained environmental conditions and so cannot build up abundance. Therefore, more species are required in order to buffer the wide range of environmental conditions experienced through time. As with spatial heterogeneity, the rate at which *b_i_* increases depends on the degree of temporal environmental autocorrelation, with the fastest increases when environmental autocorrelation is low (i.e. fastest when γ = 0, slowest when γ = 2). However, for temporal heterogeneity, *b_i_* saturates at higher levels when the environment is random compared to when it is autocorrelated. This is due to the same effect of temporal environmental heterogeneity in species performance that impacts the local scale *b_i_*.

## Discussion

We have found that the relationship between biodiversity (here the number of species) and ecosystem functioning (here total community biomass) persists at large spatial scales, but that the strength of the relationship changes with spatial and temporal scale when the environment varies. We have shown that the autocorrelation of environmental heterogeneity defines the strength of scale dependence in the biodiversity ecosystem functioning relationship (BEF). Simply put, a greater number of species is required to reach the same level of ecosystem functioning over larger spatial and temporal scales if there is more environmental heterogeneity at those larger scales. [2] Indeed, the degree of autocorrelation in environmental conditions determines how quickly this species turnover occurs and thus how quickly the number of species required to maintain ecosystem functioning increases with scale. Thus, the scale dependence in the BEF relationship is driven by the rate of species turnover that is required to maintain functioning across locations that differ in environment conditions [2]. Further, the number of species required to maintain function increases fastest with scale when environments have low autocorrelation because a wide range of conditions are encountered over relatively short spatial or temporal scales.

In our model, complementarity and selection effects both contribute to the positive BEF relationship, but their relative contribution shifts with scale. At the small or short scales where species interactions play out, complementarity occurs because intraspecific competition is stronger than interspecific competition (i.e. resource complementarity) [33]. At these scales, selection effects occur because species that are well suited to the abiotic environment produce more biomass than those that are poorly suited. Over larger or longer scales, when environmental conditions vary, these selection effects switch to being spatial or temporal complementarity effects as different species contribute to functioning in different environments [36].

The degree of environmental autocorrelation is a key determinant of species turnover and thus how quickly the number and composition of species varies with spatial or temporal scale [26]. When environmental conditions are autocorrelated, conditions are similar over small spatial or temporal scales, so that the full range of environmental conditions are only found over large spatial scales. Thus, compositional turnover is low and the number of species required to maintain ecosystem functioning increases relatively slowly with scale. In contrast low autocorrelation results in fast environmental change, and so compositional turnover is fast. In general, we expect the scale of compositional turnover to be set by distance and environmental autocorrelation [26,37,38]. Differences in composition turnover between plant and animal communities is likely linked to the scales over which they move and experience environmental autocorrelation in the conditions that affect fitness [39]. These general qualitative findings are consistent when considering scaling across space or time, but some quantitative differences emerge for space versus time. Differences between results for time versus space are due to the fact that spatial and temporal environmental heterogeneity affects whether communities are at static equilibrium with respect to the environment, which we discuss in more detail below.

### Comparing BEF scaling arising from heterogeneity in space versus time

Although spatial and temporal environmental heterogeneity both result in similar changes in the BEF relationship with scale, they have different impacts on community turnover, which has implications for how we can detect the contributions of space and time to the scaling of BEF. This is most evident over small and short scales (Fig. 2). These differences between space and time arise because of the way that spatial versus environmental changes affect whether communities are at static equilibrium with respect to environmental conditions.

The composition of a community at a given time depends greatly on its previous composition, and so communities are unlikely to be at static equilibrium with respect to a temporally varying environment. When environmental change is fast (i.e. autocorrelation is low; Fig. 1E) communities tend to track the averaged conditions. Thus, communities contain species at lower abundances because the environment is rarely ideal for them (Fig. 2D), and the species that do persist are those that are suited to average environmental conditions. When conditions change more slowly (i.e. autocorrelation is high; Fig. 1F), the community composition tracks environmental conditions but ‘lags’ behind because there is compositional inertia [15]. In contrast, when the environment is constant through time, but varies across space, each site is at equilibrium with respect to its local environment, regardless of the degree of autocorrelation (Fig. 1A-C).

In addition to driving different rates of biodiversity turnover, spatial and temporal environmental heterogeneity also impact the BEF relationship in different ways because they affect the levels of ecosystem functioning that are maintained at a given level of species diversity. By holding communities away from their static equilibrium, temporal environmental heterogeneity decreases the average biomass that is maintained in the communities compared to when heterogeneity is across space (Fig. 3). Previous research noted this and found that temporally autocorrelated environments generate low-frequency and time-lagged fluctuations in biomass that reduce the contribution of diversity to biomass production at least over short temporal scales [15].

The outcome of these differential patterns of diversity and functioning mean that the degree of environmental autocorrelation has larger impacts in temporally varying environments as compared to the spatial scenarios (Fig. 4). Whereas the level of biodiversity required to maintain functioning—at either the smallest or largest scales—is similar across all types of spatial environmental heterogeneity (Fig. 4A), this is not the case for temporal heterogeneity. With temporal environmental heterogeneity, the number of species required to maintain functioning decreases as autocorrelation increases (Fig. 4B). This is because more species are required to maintain functioning in our simulations when environmental conditions change rapidly.

Our findings differ from expectations from previous simulations that found that compositional turnover should cause the slope of the BEF relationship to increase with scale [3]. In fact, the increase in half saturation constant with scale in the present study actually corresponds to a decrease in BEF slope with scale if the saturation level of functioning remains constant. The reason for this difference is the mechanism responsible for composition turnover in the two studies. Here, turnover is due to environmental variation, which causes species to vary in abundance across space or time. In Thompson *et al*. [3] species turnover occurs through stochasticity and all species contribute equally whenever they are present. Taken together, these two approaches highlight the general expectation that compositional turnover should lead to a scale dependent BEF relationship, and whether this compositional turnover is driven by environmental factors or stochastic processes should determine if the BEF slope increases or decreases with scale.

### Testing this theory

This work provides theoretical predictions that can be tested empirically, in the lab with microcosm experiments, or in controlled surveys and experiments in the field where environmental gradients can be modified or controlled over extended spatial and temporal scales. At larger scales, remote sensing data can be used to link processes, such as primary production (on land or in the oceans), to turnover in taxonomic and functional diversity caused by shifting spatial and temporal environmental gradients obtained from global weather networks. Paleodata may also present an opportunity to assess BEF scaling over thousands of years of historical change in community composition [40]. Each of these approaches offers a different way to test our theoretical predictions, either by exploiting existing environmental gradients or manipulating them to change the scales over which environments are experienced by different assemblages.

### Model assumptions and caveats

Our sensitivity analyses (Appendix 2) show that our main conclusion—that environmental heterogeneity should lead to a greater number of species being required to maintain a given level of ecosystem functioning at larger and longer scales—is robust to the specific assumptions that we have made in our model. However, there are three assumptions to which our findings are likely sensitive. The first is that species differ in the environmental conditions in which they are most productive. Without this condition, environmental heterogeneity would not result in community turnover. The second is that species contribute to ecosystem functioning in proportion to their abundance or biomass. This is key, as environmental heterogeneity drives variation in abundance in our model, and thus functioning. The third assumption is that dispersal is not so high as to homogenize communities across space. If dispersal is so high as to homogenize communities across space, then the diversity of the community would be constant across scales and the BEF relationship would be scale invariant [13,20]. We discuss the implications of these assumptions in greater detail in our supplementary discussion (Appendix 3).

### Conclusions

Overall, this research formalizes the theoretical expectation that environmental heterogeneity should cause the relationship between biodiversity and ecosystem functioning to depend on the spatial or temporal scale at which it is observed. Our results suggest that at small scales, relatively low levels of biodiversity can sustain ecosystem functioning, but that we need to preserve biodiversity because high levels are necessary to maintain functioning over larger and longer scales where environmental heterogeneity is greater. Because environmental heterogeneity drives the BEF relationship, our results also demonstrate that the relationship between BEF and scale depends on the degree of autocorrelation of environmental conditions. This theory can help us to predict how human activities—which are changing the composition and diversity of ecological communities [41], as well the range, heterogeneity, and autocorrelation of environmental conditions that they experience [42,43]—should impact the relationship between biodiversity and ecosystem functioning from local to global scales [6].

## Acknowledgements

We thank all members of the BEF Scale working group for valuable discussions and feedback. In particular we thank Diane Srivastava for her contributions to the design of the study. This paper benefited from discussions held in the GDR TheoMoDive.

## Funding

PLT was supported by NSERC and the Killam Trust. SK was supported by the ANR project ECONET (ANR-18-CE02-0010). AG acknowledges the support of NSERC, Killam Fellowship, the Quebec Centre for Biodiversity Science (QCBS) and the Liber Ero Chair in Biodiversity Conservation. YRZ, CdM and ML were supported by the TULIP Laboratory of Excellence (ANR-10-LABX-41), and by the BIOSTASES Advanced Grant, funded by the European Research Council under the European Union’s Horizon 2020 research and innovation programme (grant agreement no. 666971). SW was supported by the National Natural Science Foundation of China (31988102). This paper arose from a joint working group supported by the Quebec Centre for Biodiversity Science (working group #12) and the Centre for Biodiversity Theory and Modelling (Moulis, France).

## Notes

### Competing Interest Statement

The authors have declared no competing interest.

### Summary of Updates

This version of the manuscript has been revised following peer review. No quantitative changes have been made to the results or findings.

