## Supplemental materials for "Scaling up biodiversity ecosystem functioning relationships: the role of environmental heterogeneity in space and time"

### Appendix 1

### Supplementary methods

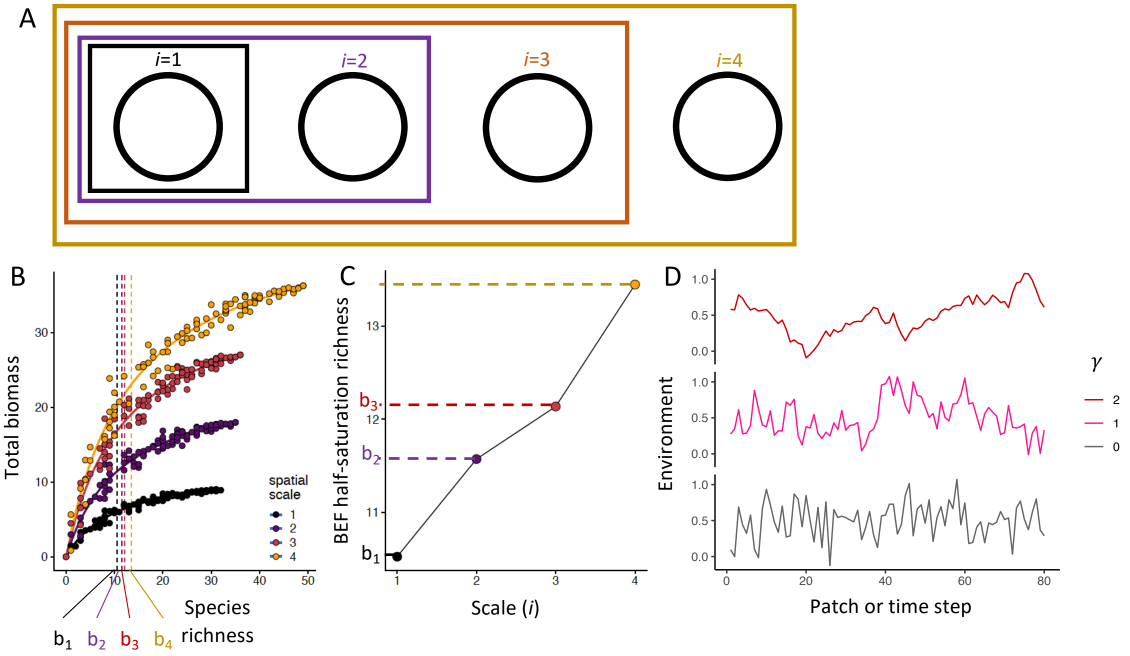


**Figure S1.** We investigated how the form of the BEF relationship (i.e., slope and asymptote) changes depending on the spatial and temporal scale at which it is measured. We used a model governed by Lotka-Volterra dynamics. In panel A, each black circle represents a community (or 'patch'), at different spatial locations or moments in time. For each community, we fit a Michaelis-Menten function to the relation between average biomass and species richness to estimate the half saturation constant (B), i.e. the number of species required to produce 50% of the biomass (parameter $b_i$). The four curves represent BEF relationships at different scales. The black curve is the BEF obtained in a single patch at a single moment (i.e. scale = 1). The other curves are obtained by aggregating data from *N* neighbouring communities (or time points for temporal scenarios). In panel C, we then plot the change in $b_i$—the half saturation of the BEF relationship— as a function of scale. We repeated this procedure for three different levels of environmental autocorrelation (see example environmental sequences in panel D; γ = 0, 1, 2 respectively), which we hypothesized to govern the rate of turnover of species richness through time or across communities. This procedure allowed us to assess how the slope and asymptote of the scaling relationship changed with environmental autocorrelation.

*Rationale for decisions in simulation*

The extinction threshold of 0.05 is necessary to ensure that species can be lost from the communities. Otherwise, biomass levels would decrease to extremely small values but never reach zero. The reintroduction step allows species to colonize the community following environmental change and ensures that we obtain temporal turnover in composition rather than just an erosion of diversity when the environment changes. This is equivalent to a low rate of dispersal from an external species pool. Because the reseeded biomass is below the extinction threshold and occurs after we record the occurrences and biomass, species will not show up in the community unless they increase above the extinction threshold within a single time step. This recolonization is not needed in the spatial scenario because the environment is constant through time, but we include it to keep the scenarios comparable.

The specific parameter values used in our model were chosen to ensure that: 1) there is turnover in community composition in response to environmental heterogeneity (i.e. $\sigma$ = 0.25); 2) that species are capable of a stable coexistence within a local patch if they are suited to the local environmental condition (i.e. interspecific competition is weaker than intraspecific competition; $\alpha_{ji} < \alpha_{ii}$); 3) when experiencing optimal environmental conditions, species have an equilibrium abundance that far exceeds the extinction threshold (i.e. $r_{max}/(1+\alpha_{ii}) > 0.25$); 4) that the initial biomass values are below this equilibrium value but above the extinction threshold. However, our results are robust to variation of all of these key parameters, provided these conditions are still met (see sensitivity analysis below).

*Generating the temporal burn-in sequence*

A shorter burn-in was used of the temporal scenario (80 time steps) than in the spatial scenario (150 time steps) because there was no need to give the community time to reach a static equilibrium, which is only reached when the environment is held constant (Figure S2). We generated this temporal burn-in sequence as a second time series rather than extending the length of the initially generated time series because the level of autocorrelation of a time series depends on its length. Thus, if we had generated a single 160 time step series with $\gamma = 2$, then $\gamma \neq 2$ for the sampled final 80 time steps. The burn-in sequence was generated in the same way as the sampled time series, but for 81 time steps. We then rescaled the burn-in time series so that its value at time step 81 was equal to the first time step of the sampled time series. This was necessary to ensure that there was no large environmental jump after the burn-in period. We then removed time step 81 of the burn-in sequence to avoid the repeated value in the final time series. To ensure that the range of conditions in the burn-in series were similar to the sampled time series, we excluded any time series which contained values greater than 1.3 or less than -0.3.

*Static and Dynamic equilibria*

While the communities in our spatial scenario simulations are at static equilibrium with respect to the environmental conditions, the communities in the temporal simulations are not. That is, their composition in a given environment is not the same as it would be if the environment were held constant. This is evident in Figure 1, where the species present have optima that are close to the environmental conditions (the red line) in the spatial scenarios (Fig. 1A-C) but have optima that are further from the environmental conditions when the environment varies through time (Fig. 1D-F). The static equilibria in the spatial scenarios is also evident in Fig. S2A-C, which shows how species richness changes throughout the burn-in period when all species are included in the simulation (i.e., $S^* = 100$). Here, the species richness in each patch decreases from the initial 100 species and reaches an equilibrium around 30 species. A similar equilibrium is reached in the temporally varying scenarios (Fig. S2D-F), but in this case it is a dynamic equilibrium. That is, the number of species is relatively stable through time, but it is a different number than when the environment is held constant. This number of species that are maintained during the temporally varying environment fluctuate around this dynamic equilibrium but extending the length of the time series would not stabilize these fluctuations or change their value.

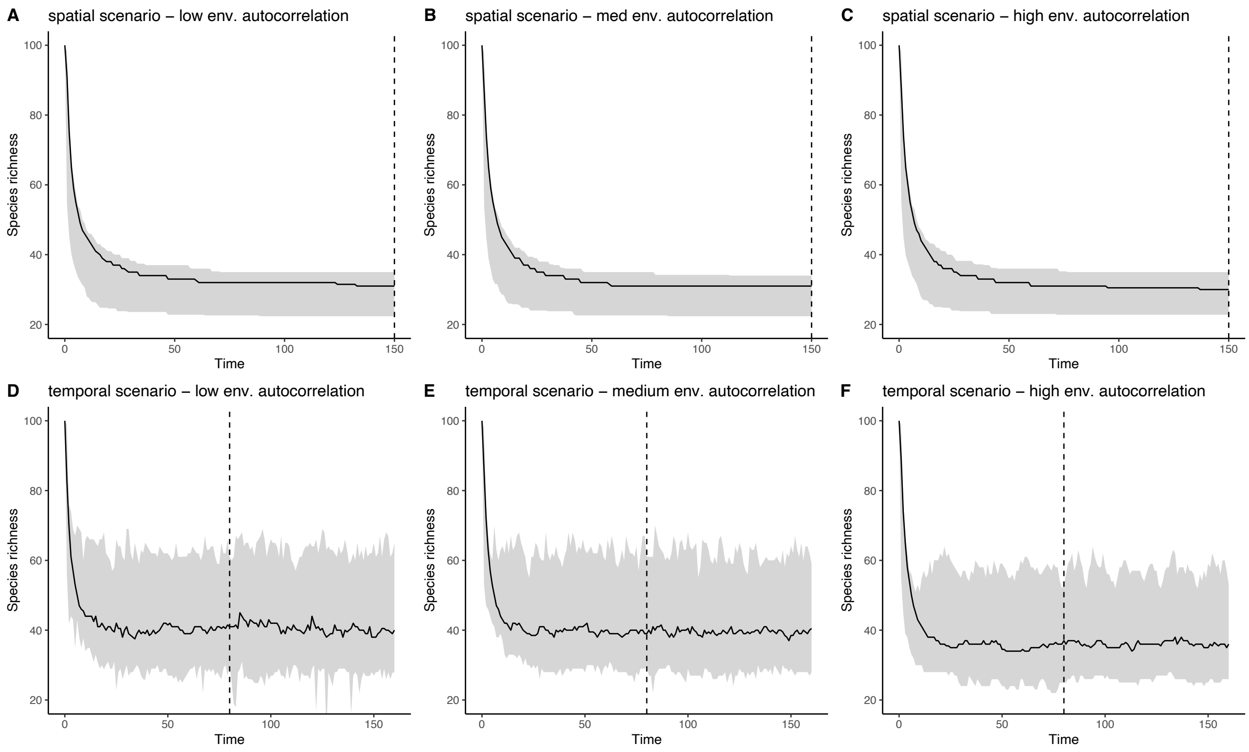


**Figure S2.** The number of species present in an individual habitat patch throughout the burn-in period (before the vertical dashed line) and during the period considered in the main analysis (time step 150 for the spatial scenarios and time steps 81-160 in the temporal scenarios). Panels A – C show the three spatial scenarios, where the environment in each patch is held constant the entire time. Panels D – F show the three temporal scenarios, where the environment in each patch varies through time. Low, medium, and high environmental autocorrelation are shown in panels A and D, B and E, and C and F, respectively. The line shows the median value across 100 replicate patches, the shaded area shows the 1-99% quantile range.

**Appendix 2**

**Sensitivity analyses**

We performed a series of analyses of key parameters to test the sensitivity of our results to the values of these parameters. When testing the sensitivity of each parameter, we held all other parameters at the default value used in the main analysis. We then ran 20 replicates of the simulation where we decreased the value of the parameter (Figure S3) and another 20 replicates where we increased the value of the parameter (Figure S4). The parameter values used are outlined in Table S1. In all cases, changing the values of the parameters did not change the qualitative patterns, compared to the main analysis.

We also performed an analysis to test the sensitivity of our findings to the assumption of equal intraspecific competition across all species. In this case, we drew values of the intraspecific competition coefficient $\alpha_{ii}$ for each species from a normal distribution with a mean of 1 (corresponding to the value used in our main analysis) and a standard deviation of 0.25. We found that interspecific variation in the strength of intraspecific competition did not change the qualitative patterns compared to the main analysis (Figure S5).

**Table S1.** Parameters tested in the sensitivity analyses, their interpretations and the values used in the analyses. The value used in the main simulations is underlined.

| Parameter | Meaning | Values tested |
|---|---|---|
| Maximum $\alpha_{ij}$ | The upper limit of the range from which interspecific competition values are drawn. The lower limit is 0. | 0.1, <u>0.25</u>, 0.5 |
| max scale | The maximum spatial or temporal scale | 40, <u>80</u>, 160 |
| $r_{max}$ | The maximum density-independent growth rate, which occurs when the local environmental conditions match the species environmental optimum | 3, <u>5</u>, 10 |
| $\sigma$ | The width of the environmental niche | 0.2, <u>0.25</u>, 0.3 |

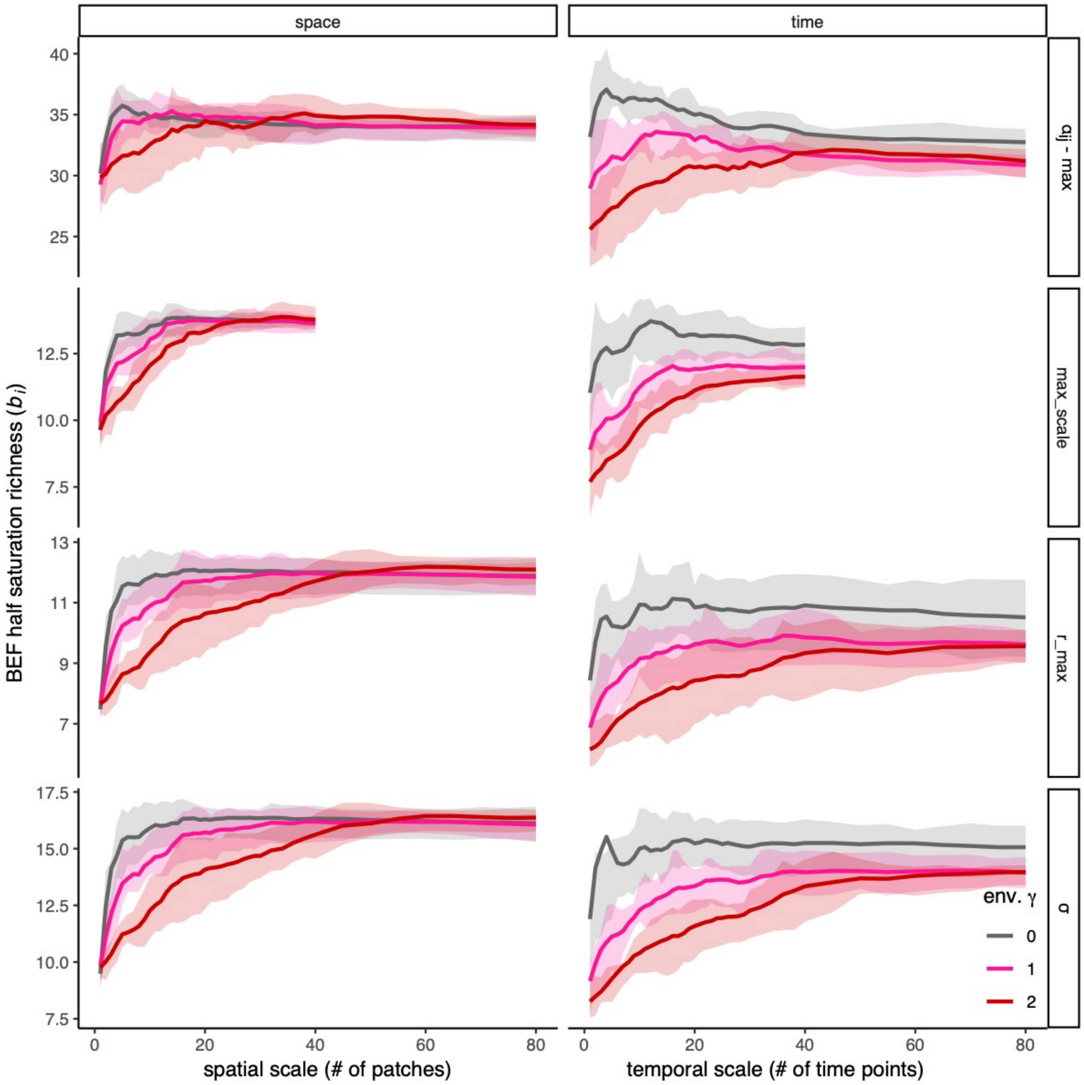


**Figure S3.** Sensitivity analysis showing results from 20 replicate runs where a single focal parameter was decreased to level lowest value in Table S1. The figure shows the relationship between the BEF half saturation richness $b_i$ and the spatial (left column) or temporal (right column) scale. The rows correspond to the four parameters used in the sensitivity analysis as indicated by the labels on the right side. In each case, all parameters other than the focal parameter were set to the default value used in the main analysis.

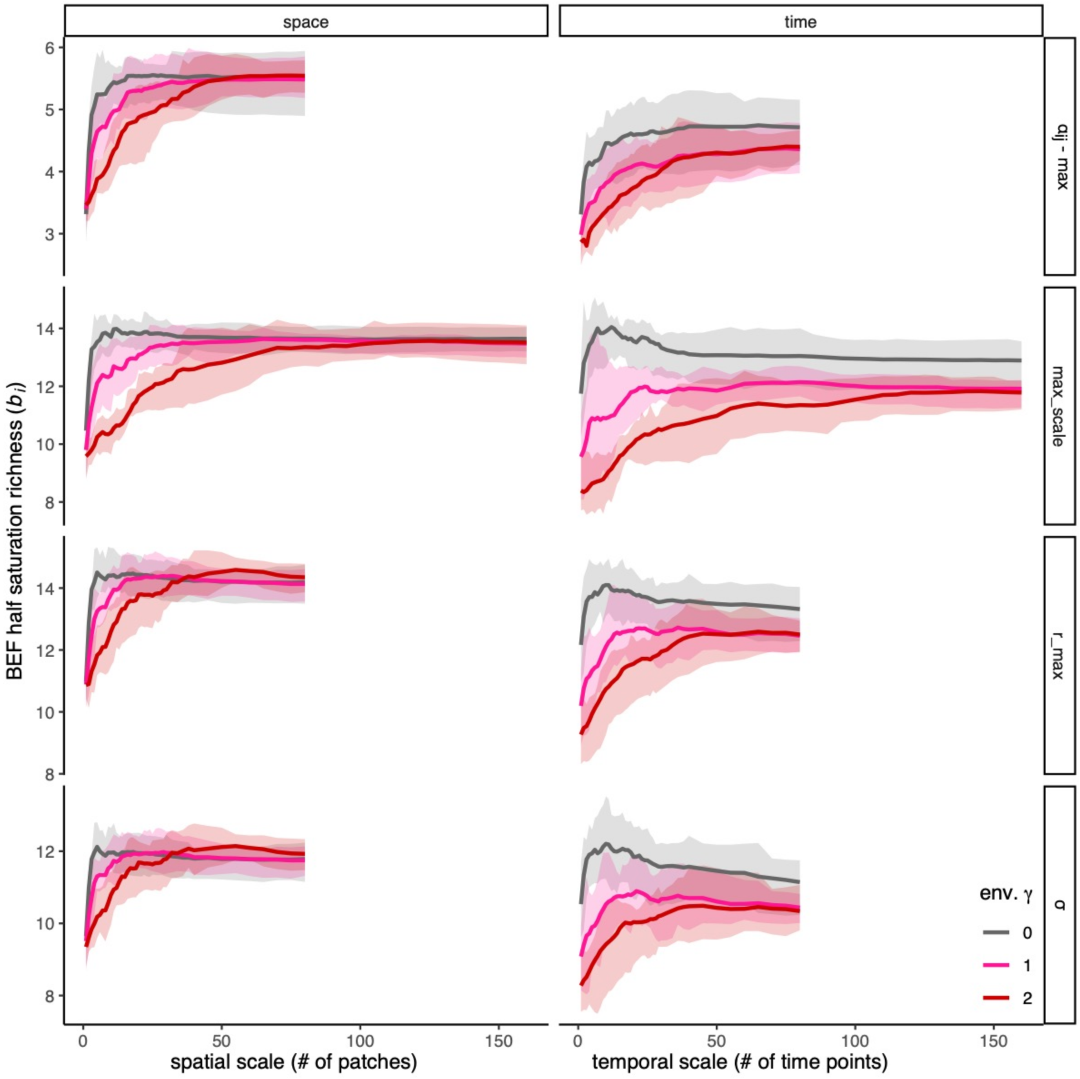


**Figure S4.** Sensitivity analysis showing results from 20 replicate runs where a single focal parameter was increased to highest value level in Table S1. The figure shows the relationship between the BEF half saturation richness $b_i$ and the spatial (left column) or temporal (right column) scale. The rows correspond to the four parameters used in the sensitivity analysis as indicated by the labels on the right side. In each case, all parameters other than the focal parameter were set to the default value used in the main analysis.

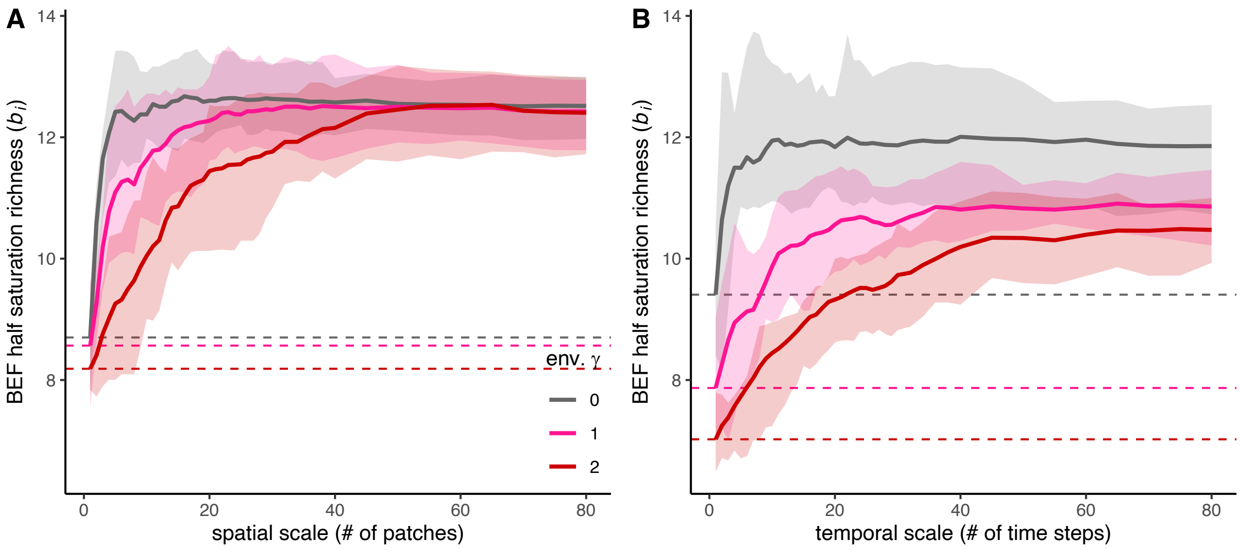


**Figure S5.** Sensitivity analysis showing the relationship between the half saturation species richness of the BEF relationship and the spatial (A) or temporal (B) scale of observation, when intraspecific competition is allowed to vary across species. Colour indicates the γ value of the degree of autocorrelation of environmental heterogeneity. Dashed lines show the half saturation species richness at a scale of 1. Lines represent the mean and bands represent the interquartile range across 20 replicate simulations.

**Appendix 3**

**Supplementary discussion**

*Model assumptions and caveats*

We expect that our main conclusion—that environmental heterogeneity should lead to a greater number of species being required to maintain a given level of ecosystem functioning at larger and longer scales— is most dependent on our assumption that species differ in the environmental conditions in which they are most productive. As discussed in the main manuscript, beta diversity that results from demographic stochasticity rather than environmental heterogeneity may actually have the opposite effect. A valuable next step would be to compare the scale dependence of the BEF in a single model while varying species sensitivity to environmental heterogeneity [sensu 1,2]. Because the scale dependence of the BEF relationship that we find here is driven by environmental heterogeneity, we do not expect our findings to depend strongly on our assumptions about the strength of competition between species. Although positive BEF relationships at small/short scales depend on the assumption that intraspecific competition is stronger than interspecific competition, this is not the case at larger/longer scales, where complementarity can arise through differential responses to spatial environmental heterogeneity [3].

We have assumed that ecosystem functioning is proportional to the abundance of species in our communities. This is likely to be a reasonable assumption when individuals of different species are of comparable size, and when considering ecosystem properties such as biomass. However, in cases when rare species contribute disproportionately to functioning [4,5], our findings may be less relevant. However, as long as environmental heterogeneity results in spatial and temporal variation in species performance, our conclusion that more species are required to maintain functioning over longer and larger scales should still hold.

Finally, we do not expect our main conclusion to be greatly influenced by our assumptions about dispersal. But we do expect that dispersal should affect the degree to which spatial and temporal environmental heterogeneity differ in how they cause the BEF relationship to change with scale. High rates of dispersal between nearby sites would produce mass effects and would cause communities to homogenize [6]. With extremely high rates of dispersal, the spatial and temporal patterns should converge, but this would require dispersal of more than half of the individuals at each time step, after which the temporal inertia of the communities would be overwhelmed. In addition, we expect that such spatial homogenization would reduce the differences between levels of spatial environmental autocorrelation.
